# The role of transposable elements for gene expression in *Capsella* hybrids and allopolyploids

**DOI:** 10.1101/044016

**Authors:** Kim A. Steige, Johan Reimegård, Carolin A. Rebernig, Claudia Köhler, Douglas G. Scofield, Tanja Slotte

## Abstract

The formation of an allopolyploid species involves the merger of two genomes with separate evolutionary histories. In allopolyploids, genes derived from one progenitor species are often expressed at higher levels than those from the other progenitor. It has been suggested that this could be due to differences in transposable element (TE) content among progenitors, as silencing of TEs can affect expression of nearby genes. Here, we examine the role of TEs for expression biases in the widespread allotetraploid *Capsella bursa-pastoris* and in diploid F1 hybrids generated by crossing *Capsella orientalis* and *Capsella rubella*, two close relatives of the progenitors of *C. bursa-pastoris*. As *C. rubella* harbors more TEs than *C. orientalis*, we expect *C. orientalis* alleles to be expressed at higher levels if TE content is key for expression biases. To test this hypothesis, we quantified expression biases at approximately 5800 genes in flower buds and leaves, while correcting for read mapping biases using genomic data. While three of four *C. bursa-pastoris* accessions exhibited a shift toward higher relative expression of *C. orientalis* alleles, the fourth *C. bursa-pastoris* accession had the opposite direction of expression bias, as did diploid F1 hybrids. Associations between TE polymorphism and expression bias were weak, and the effect of TEs on expression bias was small. These results suggest that differences in TE content alone cannot fully explain expression biases in these species. Future studies should investigate the role of differences in TE silencing efficacy, as well as a broader set of other factors. Our results are important for a more general understanding of the role of TEs for cis-regulatory evolution in plants.

## Introduction

Polyploidy, or whole genome duplication (WGD), is a key contributor to plant speciation. It has been estimated that about 15% of speciation processes in angiosperms involve polyploidy (Wood et al 2009), and most flowering plant species have experienced WGD at some point during their history. Information on the genomic consequences of WGD is therefore important for understanding plant genome evolution. Polyploids can form either through WGD within one species (autopolyploidy) or by hybridization and genome doubling (allopolyploidy) (Ramsey and Schemske 1998). The formation of an allopolyploid species thus involves the merger of two genomes with separate evolutionary histories, a fact that is thought to have a major impact on establishment and persistence (Soltis et al 2009; Barker et al 2015) and on subsequent genome evolution in allopolyploids (Freeling et al 2012; Steige and Slotte 2016).

A common observation in both recent and ancient allopolyploids is that homeologous genes on one subgenome tend to be expressed at a higher level than those on the other subgenome (e.g. *Gossypium;* Flagel and Wendel 2010, *Brassica*; Woodhouse et al 2014, wheat; Li et al 2014, maize; Schnable et al 2011). Such systematic homeolog expression biases are thought to affect the evolutionary trajectories of duplicate genes, ultimately leading to patterns of biased fractionation (Langham et al 2004) through preferential loss of the less expressed copy (Schnable et al 2011). Understanding how patterns of systematic homeolog expression bias are established is therefore of general importance for understanding processes that govern genome evolution in allopolyploids.

It has recently been suggested that differences between the progenitor species in their content of transposable elements (TEs) might be key for the establishment of homeolog expression bias (Schnable et al 2011; Freeling 2012; Woodhouse et al 2014). This is thought to occur at least in part through the RNA-directed DNA methylation (RdDM) pathway, a major pathway for silencing of TEs in plants (Matzke and Mosher 2014). It is known that silencing of TEs through the RdDM pathway can also result in repression of nearby genes (Lippman et al 2004; Hollister and Gaut 2009; Hollister et al 2011), and there is evidence for a role of TE silencing for cis-regulatory variation in *A. thaliana* (Wang et al 2013) and for *cis-* regulatory divergence among crucifers (Hollister et al 2011; Steige et al 2015a). Recently, it was suggested that silencing of TEs through the RdDM pathway might also be of general importance for establishing patterns of homeolog expression bias in allopolyploids (Freeling et al 2012). Under this model, silencing of TEs results in preferential expression of genes present on the subgenome that harbors fewer TEs in the vicinity of genes. In allopolyploids, as well as in diploid hybrids, we would thus expect to see a higher relative expression of genes from the progenitor that has a lower content of TEs near genes, and/or a lower efficacy of silencing of TEs.

The crucifer genus *Capsella* is a suitable system for testing for a role of TEs in establishing homeolog expression biases. The *Capsella* genus contains three diploid species (Chater et al 1993); the outcrosser *Capsella grandiflora* and the selfers *Capsella orientalis* and *Capsella rubella*, as well as the tetraploid *Capsella bursa-pastoris*, which has a nearly worldwide distribution as a highly successful weed (Hurka and Neuffer 1997). *C. bursa*-pastoris is a highly self-fertilizing tetraploid with disomic inheritance, i.e. the two homeologous subgenomes are independently inherited. We have recently shown that *C. bursa-pastoris* formed within the last 300 ky by hybridization and genome duplication involving two ancestral species, *C. orientalis* and an ancestor of the diploids *C. grandiflora* and *C. rubella* (Douglas et al. 2015). The two progenitors of *C. bursa-pastoris* likely differed in TE content, as the genomes of *C. grandiflora* and *C. rubella* both have a markedly higher TE content than *C. orientalis*, both genome-wide and near genes (Ågren et al 2014), and *C. bursa-pastoris* does not appear to have undergone large-scale proliferation of TEs since its origin (Ågren et al 2016). Thus, we would expect to see a global shift toward higher expression of alleles derived from *C. orientalis* in the allopolyploid *C. bursa-pastoris*. Under this model, we would also expect to see the same shift in diploid hybrids derived from the closest extant relatives of the progenitors of *C. bursa-pastoris*. However additional genome-level features may act counter to these expectations, for example, if the two progenitors of *C. bursa-pastoris* differ in their efficacy of silencing TEs, or if polyploidy has additional effects on gene expression.

Here, we tested these hypotheses by assessing homeolog-specific expression (HSE) in the tetraploid *C. bursa-pastoris* and allele-specific expression (ASE) in diploid hybrids generated by crossing two diploid close relatives of the progenitors of *C. bursa-pastoris*, namely *C. rubella* and *C. orientalis*. We generated deep transcriptome and genomic sequencing data and mapped against parental haplotypes to reduce mapping bias. Deep transcriptome sequencing data was analyzed to quantify expression biases in flower buds and leaves of both natural allopolyploids and diploid hybrids, while correcting for technical variation and read mapping biases using genomic data. This allowed us to test for a directional shift in expression of alleles from each progenitor lineage and to test whether expression biases are associated with TE insertions near genes. Finally, we assessed to what extent homeolog expression biases in allopolyploids reflected *cis*-regulatory divergence among the diploid progenitors. Our results are important for an improved understanding of the processes that govern expression evolution upon hybridization and allopolyploidization in plants.

## Results

### Sequencing data and processing

We generated transcriptome and genome sequencing data for four accessions of the allopolyploid *C. bursa-pastoris* (Table S1), representing the major genetic clusters identified in this species (Slotte et al 2009; Cornille et al 2016), and two diploid F1 hybrids derived from crosses of *C. orientalis* and *C. rubella* (Table S2). In total, we obtained 317.6 Gbp of high-quality (Q≥30) RNAseq data from flower buds and leaves, and for each accession or cross, we included three biological replicates (Table S3). To account for effects of technical variation on allele-specific expression and to reconstruct parental haplotypes, we further generated whole genome resequencing data (total 80.2 Gbp Q≥30, expected mean coverage 32x) for all included individuals as well as the *C. orientalis* and *C. rubella* parents of the diploid F1 hybrids (Table S4).

We took several steps to avoid effects of read mapping artifacts in our analyses of ASE in the diploid hybrids, and HSE in the tetraploids. First, we mapped RNAseq data to parental haplotypes reconstructed based on genomic data, and conducted stringent filtering to avoid inclusion of low-confidence SNPs, as in Steige et al (2015a) (see Methods for details). We further identified a set of coding SNPs where we could confidently assign each allele in *C. bursa-pastoris* to the A homeolog derived from the *C. grandiflora/C. rubella* lineage (CbpA), or the B homeolog derived from *C. orientalis* (CbpB) (see Methods for details). After these filtering steps, we retained approximately 5770 genes with 27680 transcribed SNPs that were amenable for analyzing transcriptomic biases (Table 1). The median allelic ratio of genomic read counts at these SNPs was 0.504 (range 0.492 to 0.516), suggesting that there was little remaining mapping bias in our data (see Figures 1 - 4). Our bioinformatic procedures also greatly reduced mapping bias compared to results when mapping to the *C. rubella* reference genome (Supplementary Figure S1 & S2).

**Table 1.** Genes amenable to analysis of ASE in flower bud and leaf samples from the two diploid *C. orientalis* x *C. rubella* F1s and the four *C. bursa-pastoris* accessions, counts of genes with evidence for ASE and the estimated false discovery rate (FDR) and proportion of genes with ASE.

| Sample designation | Type | Sample | Genes amenable to ASE analysis<sup>1</sup> | Analyzed genes<sup>2</sup> | Heterozygous SNPs in analyzed genes | Number of genes with ASE PP ≥ 0.95<sup>3</sup> | FDR | ASE proportion<sup>4</sup> |
| --- | --- | --- | --- | --- | --- | --- | --- | --- |
| Inter13 | diploid F1 | Flower buds | 5850 | 5739 | 28476 | 1812 | 0.00368 | 0.492 |
| Inter14 | diploid F1 |  | 5635 | 5547 | 27089 | 1995 | 0.00323 | 0.534 |
| CbpDE | *C. bursa-pastoris* |  | 5729 | 5633 | 27990 | 2240 | 0.00324 | 0.572 |
| CbpGR | *C. bursa-pastoris* |  | 5806 | 5705 | 28323 | 2019 | 0.00280 | 0.519 |
| CbpKMB | *C. bursa-pastoris* |  | 5814 | 5721 | 28420 | 2847 | 0.00321 | 0.670 |
| CbpGY | *C. bursa-pastoris* |  | 5777 | 5670 | 28177 | 2554 | 0.00287 | 0.632 |
| Inter13 | diploid F1 | Leaves | 5850 | 5740 | 28482 | 1565 | 0.00258 | 0.416 |
| Inter14 | diploid F1 |  | 5635 | 5297 | 26107 | 1373 | 0.00259 | 0.398 |
| CbpDE | *C. bursa-pastoris* |  | 5729 | 5409 | 27097 | 2344 | 0.00302 | 0.606 |
| CbpGR | *C. bursa-pastoris* |  | 5806 | 5427 | 27219 | 2042 | 0.00272 | 0.541 |
| CbpKMB | *C. bursa-pastoris* |  | 5814 | 5487 | 27490 | 3000 | 0.00270 | 0.714 |
| CbpGY | *C. bursa-pastoris* |  | 5777 | 5449 | 27285 | 2775 | 0.00300 | 0.689 |

<sup>1</sup>Total number of genes with heterozygous SNPs in coding regions remaining after filtering.
<sup>2</sup>Number of genes amenable to ASE analyses with expression data in at least one of the replicates of the sample.
<sup>3</sup>Genes with evidence for ASE (posterior probability ≥ 0.95).
<sup>4</sup>Direct estimate of the ASE proportion independent of significance cutoffs.

**Figure 1.**
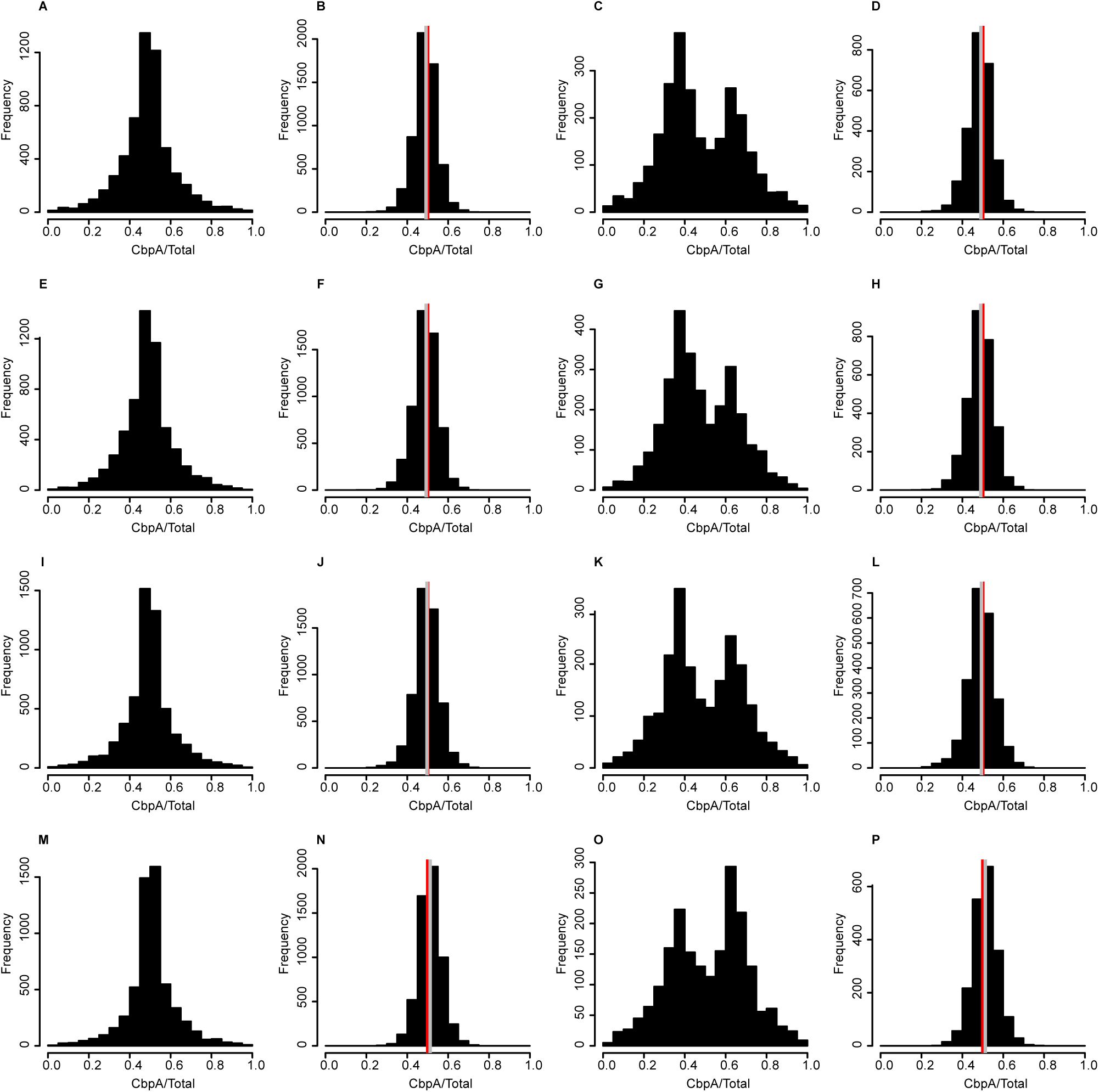
HSE in flower buds of *C. bursa-pastoris*. Histograms show the ratio of the *C. bursa-pastoris* A homeolog to total (CbpA/Total) for all expressed genes in flower buds for transcriptomic (A, E, I, M) and genomic (B, F, J, N) reads. ASE ratios for genes with significant HSE (posterior probability >= 0.95) are shown in panels C, G, K, O, and genomic ratios for the same genes are shown in panels (D, H, L, P). Genomic data have a red bar at the equal ratio (0.5) and a grey bar at the median genomic ratio plotted. Samples plotted are CbpGY (A, B, C, D), CbpKMB (E, F, G, H), CbpDE (I, J, K, L) and CbpGR (M, N, O, P).

**Figure 2.**
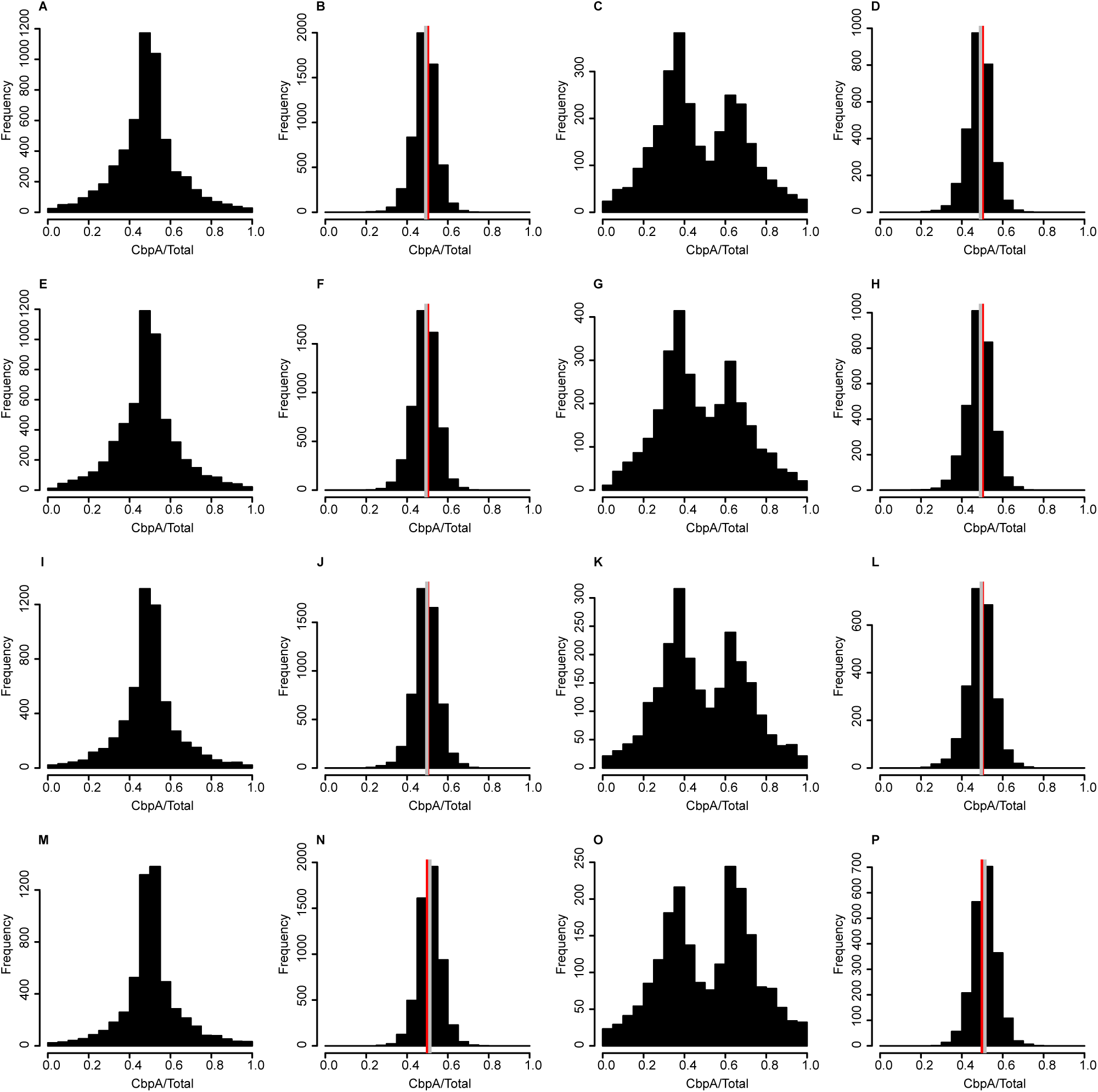
HSE in leaves of *C. bursa-pastoris*. Histograms show the ratio of the *C. bursa-pastoris* A homeolog to total (CbpA/Total) for all expressed genes in flower buds for transcriptomic (A, E, I, M) and genomic (B, F, J, N) reads. ASE ratios for genes with significant HSE (posterior probability >= 0.95) are shown in panels C, G, K, O, and genomic ratios for the same genes are shown in panels (D, H, L, P). Genomic data have a red bar at the equal ratio (0.5) and a grey bar at the median genomic ratio plotted. Samples plotted are CbpGY (A, B, C, D), CbpKMB (E, F, G, H), CbpDE (I, J, K, L) and CbpGR (M, N, O, P).

**Figure 3.**
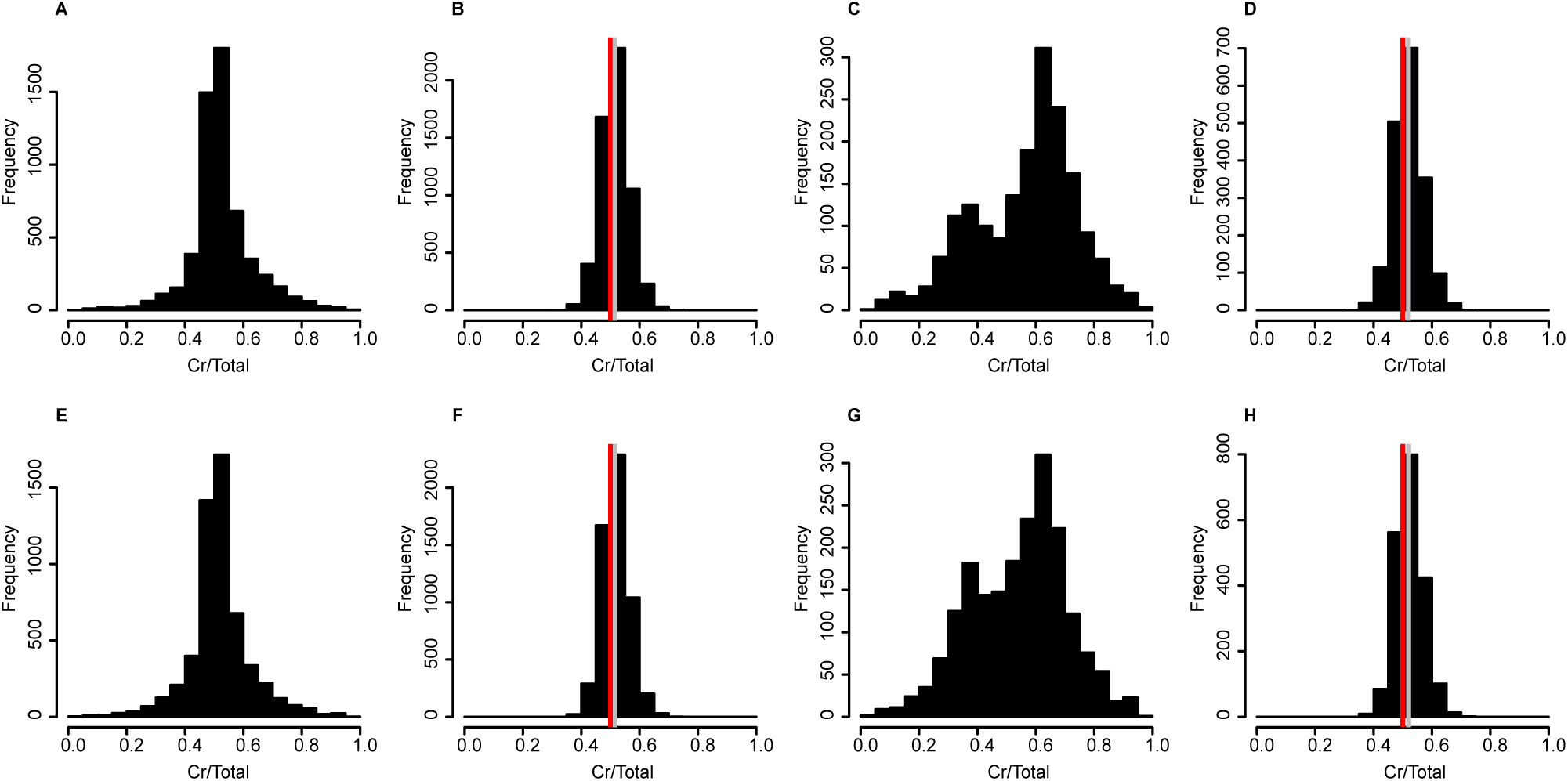
ASE in flower buds of interspecific hybrids. Ratios of the *C. rubella* allele to the total (Cr/Total) for genomic and transcriptomic data. Histograms show the distribution of ASE ratios of all expressed genes in flower buds (A, E), and for significant genes (C, G). Genomic ratios for all genes (B, F) and significant genes (D, H) are also shown. Samples plotted are Inter13 (A, B, C, D) and Inter14 (E, F, G, H). Genomic data have a red bar at the equal ratio (0.5) and a grey bar at the median genomic ratio.

**Figure 4.**
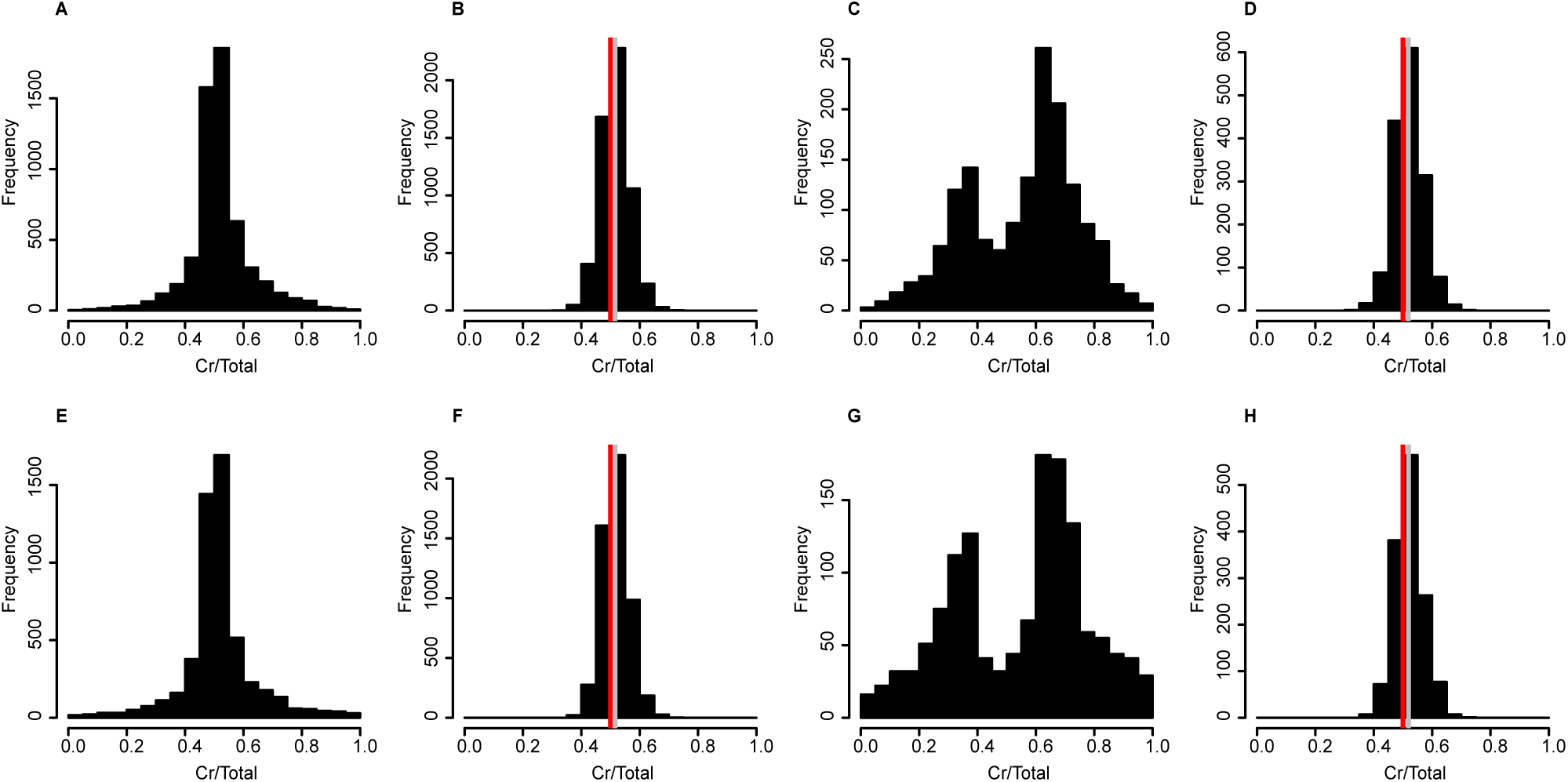
ASE in leaves of interspecific hybrids. Ratios of the *C. rubella* allele to the total (Cr/Total) for genomic and transcriptomic data. Histograms show the distribution of ASE ratios of all expressed genes in leaves (A, E), and for significant genes (C, G). Genomic ratios for all genes (B, F) and significant genes (D, H) are also shown. Samples plotted are Inter13 (A, B, C, D) and Inter14 (E, F, G, H). Genomic data have a red bar at the equal ratio (0.5) and a grey bar at the median genomic ratio.

### Homeolog expression bias in C. bursa-pastoris

For analyses of expression biases, we used a hierarchical Bayesian method developed by Skelly et al (2011), which we have previously used for analyses of *Capsella* data (Steige et al 2015a). Using this method, we estimate that a high proportion of the analyzed genes show homeolog-specific expression in *C. bursa-pastoris* (on average 59.8% vs 67.0% in flower buds and leaves, respectively, Table 1), but only 5.5% of genes showed strong expression biases (ratio of the *C. bursa-pastoris* A homeolog to total; CbpA/Total>0.8 or <0.2). The proportion of genes with HSE in *C. bursa-pastoris* was higher than the proportion of genes with ASE in *C. grandiflora* x *C. rubella* hybrids (44%; Steige et al 2015a) or within *C. grandiflora* (35%; Steige et al 2015b), as might be expected given the greater divergence of the progenitors of *C. bursa-pastoris* (Douglas et al 2015).

We next considered the direction of homeolog expression bias in *C. bursa-pastoris*. Consistent with results in Douglas et al (2015), plots of HSE for all analyzed genes show little evidence of deviation from equal expression of both homeologs (Figures 1 & 2). However, when considering only genes with a high posterior probability of HSE (PP ≥ 0.95), global shifts in the direction of HSE were evident (Figures 1 & 2). For three of the analyzed *C. bursa-pastoris* accessions, and for both flowers and leaves, there was a global shift toward higher relative expression of the B homeolog derived from *C. orientalis*. This is in agreement with expectations under a model where TE silencing is important for generating homeolog expression biases, as *C. orientalis* harbors a lower fraction of TEs both genomewide and close to genes (Ågren et al 2014; Ågren et al 2016). However, the fourth analyzed *C. bursa-pastoris* accession (CbpGR) deviated from this pattern and instead showed a bias toward elevated expression of the A homeolog (Figures 1 & 2), demonstrating that there is variation in homeolog expression bias within *C. bursa-pastoris*, and that not all accessions fit the expectations under the TE silencing model.

### Variation in homeolog-specific expression within C. bursa-pastoris

The differences among *C. bursa-pastoris* accessions in the direction of homeolog expression bias prompted us to investigate the degree of variation in HSE in *C. bursa-pastoris* (Figure 5). While all four *C. bursa-pastoris* accessions had evidence for HSE (defined as posterior probability of HSE ≥ 0.95) at a total of 1190 genes in flower buds and 1321 genes in leaves (Table 1), there were differences in the direction of homeolog expression bias among accessions for approximately half of these genes (51.9% in flower buds and 52% in leaves). Accession-specific silencing of different homeologs of the gene *FLC* in *C. bursa-pastoris* has previously been demonstrated for three of the accessions included in this study (Slotte et al 2009) and to validate our RNAseq analyses we compared our results for *FLC* to those in Slotte et al (2009). In good agreement with the results of Slotte et al (2009), our data supports a relatively strong bias toward the *FLC* A homeolog in CbpGY and CbpKMB (CbpA/Total = 0.8, posterior probability of HSE = 1 in both accessions) and a weaker but still significant bias toward the *FLC* B homeolog in CbpDE (CbpA/Total=0.32, posterior probability of HSE =1). Thus, accession-specific homeolog expression bias seems to be an important component of expression variation in *C. bursa-pastoris*.

**Figure 5.**
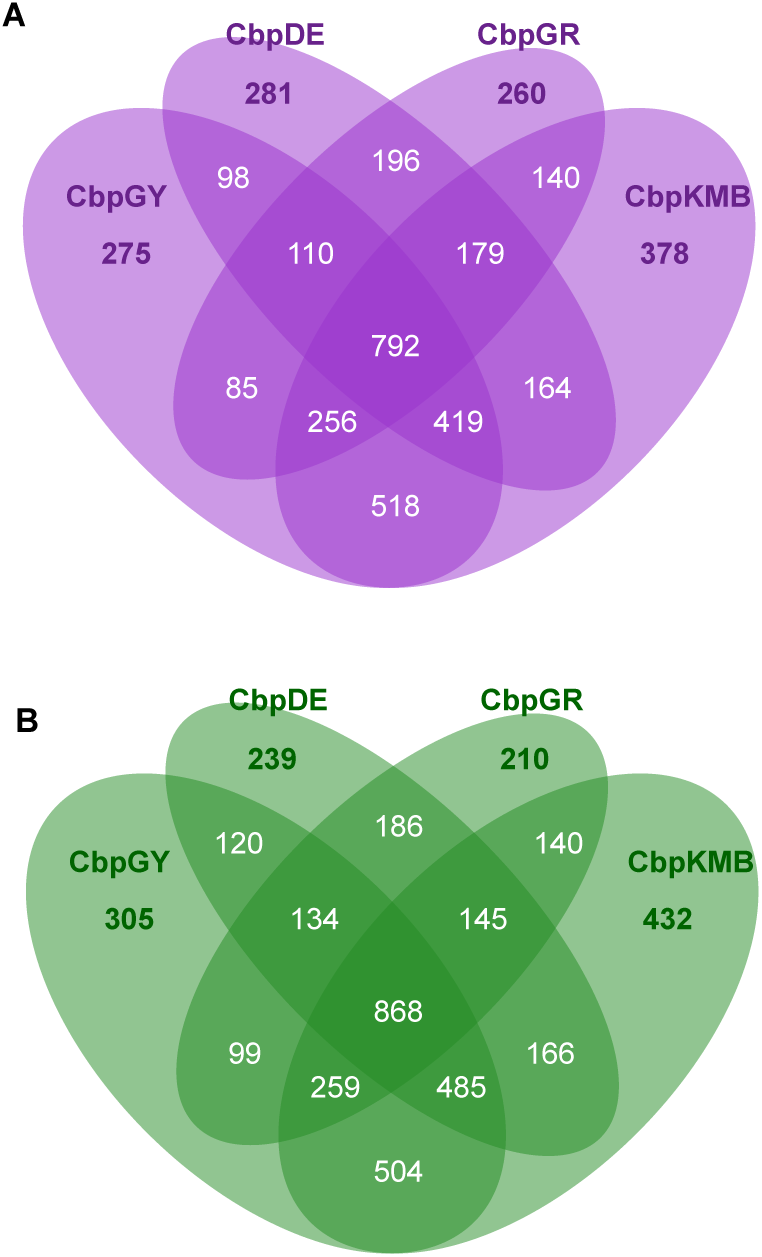
Variation of significant HSE genes in *C. bursa-pastoris*. Venn diagram for all significant genes in flower buds (A) and leaves (B).

We further considered whether there was evidence for organ-specific silencing of homeologs in *C. bursa-pastoris*. Out of a total of approximately 5417 genes that could be assessed in both leaves and flowers of *C. bursa-pastoris*, on average only 1.59% showed evidence of organ-specific silencing of one homeolog or the other, suggesting organ-specific silencing is not very frequent in the young tetraploid *C. bursa-pastoris*.

### Allele-specific expression in diploid hybrids

We next assessed allele-specific expression in the diploid F1 hybrids. In the *C. orientalis* x *C. rubella* F1s, there was evidence for ASE at a somewhat lower proportion of genes than in *C. bursa-pastoris*, 51.3% in flower buds, and 40.7% in leaves (Table 1). In the F1s, we would expect *C. orientalis* alleles to be expressed at higher levels than *C. rubella* alleles under a simple model where TE content affects *cis-*regulatory divergence as well as homeolog expression bias. Our results do not agree with this prediction. Instead, for genes with strong evidence for ASE (PP≥ 0.95) there is a shift toward lower relative expression of the *C. orientalis* allele, in both flower buds and leaves (Figures 3 & 4). Thus, the data for the diploid hybrids do not support a model where a difference in the genomewide content of TEs is the main factor underlying cis-regulatory divergence.

### Weak association between expression bias and TE insertions

To further examine the possible impact of TEs on expression divergence, we tested for an association between the presence of significant expression biases and TE insertions. We first identified TE insertions using genomic data from F1s, their *C. rubella* and *C. orientalis* parents, and tetraploid *C. bursa-pastoris* as in Ågren et al (2014) and Steige et al (2015a). Our results agree with those of Ågren et al (2014), in that we found a higher number of TE copies in *C. rubella* than in *C. orientalis* (Table 2, Table S5), and *C. bursa-pastoris* harbored slightly fewer TE insertions than the diploid F1 hybrids (Table 2, Table S6). In diploid hybrids, *Gypsy* was most abundant among heterozygous TEs, whereas *Copia* insertions were the most common among TEs called as heterozygous in *C. bursa-pastoris* (note that these likely correspond to TE insertions that differ among *C. bursa-pastoris* subgenomes, as *C. bursa-pastoris* is highly selfing and has disomic inheritance and is thus expected to be highly homozygous at each homeologous locus) (Table 2, Supplementary Table S6).

**Table 2.**
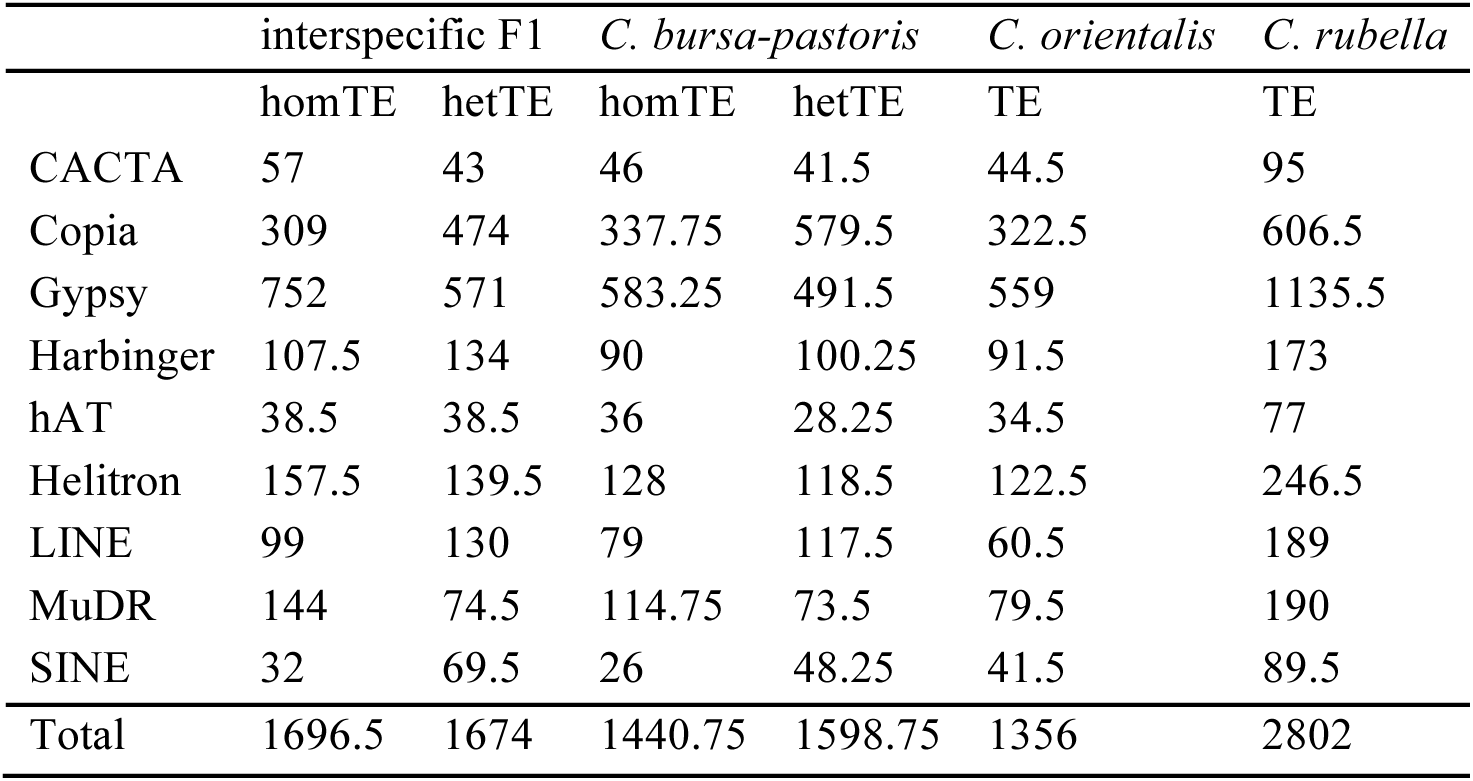
Mean abundance of Transposable Element (TE) Insertions. For interspecific *C. orientalis* x *C. rubella* F1s and *C. bursa-pastoris*, the mean number of homozygous and heterozygous TE insertions are shown (designated homTE vs hetTE, respectively).

|  | interspecific F1 |  | *C. bursa-pastoris* |  | *C. orientalis* | *C. rubella* |
| --- | --- | --- | --- | --- | --- | --- |
|  | homTE | hetTE | homTE | hetTE | TE | TE |
| CACTA | 57 | 43 | 46 | 41.5 | 44.5 | 95 |
| Copia | 309 | 474 | 337.75 | 579.5 | 322.5 | 606.5 |
| Gypsy | 752 | 571 | 583.25 | 491.5 | 559 | 1135.5 |
| Harbinger | 107.5 | 134 | 90 | 100.25 | 91.5 | 173 |
| hAT | 38.5 | 38.5 | 36 | 28.25 | 34.5 | 77 |
| Helitron | 157.5 | 139.5 | 128 | 118.5 | 122.5 | 246.5 |
| LINE | 99 | 130 | 79 | 117.5 | 60.5 | 189 |
| MuDR | 144 | 74.5 | 114.75 | 73.5 | 79.5 | 190 |
| SINE | 32 | 69.5 | 26 | 48.25 | 41.5 | 89.5 |
| Total | 1696.5 | 1674 | 1440.75 | 1598.75 | 1356 | 2802 |

While there was an association of ASE and heterozygous TEs in some diploid hybrids and *C. bursa-pastoris* accessions, patterns differed among individuals and accessions, and the strength of association was relatively weak (Figure 6). This does not seem to be a general result of reduced power due to the lower number of genes analyzed here than in Steige et al (2015a), because these results hold when analyzing a larger set of genes (~13,000 genes) in the diploid F1s (Table S7). Moreover, while there was a significant effect on heterozygous TEs on nearby gene expression in diploid F1s (Figure S3) the effect size was generally small (Figure S3, Figure S4, Table S8) and not significant after multiple testing correction. Overall, the association between expression bias and TEs was therefore weaker than that previously observed in *C. grandiflora* (Steige et al 2015b) and *C. grandiflora* x *C. rubella* hybrids (Steige et al 2015a).

**Figure 6.**
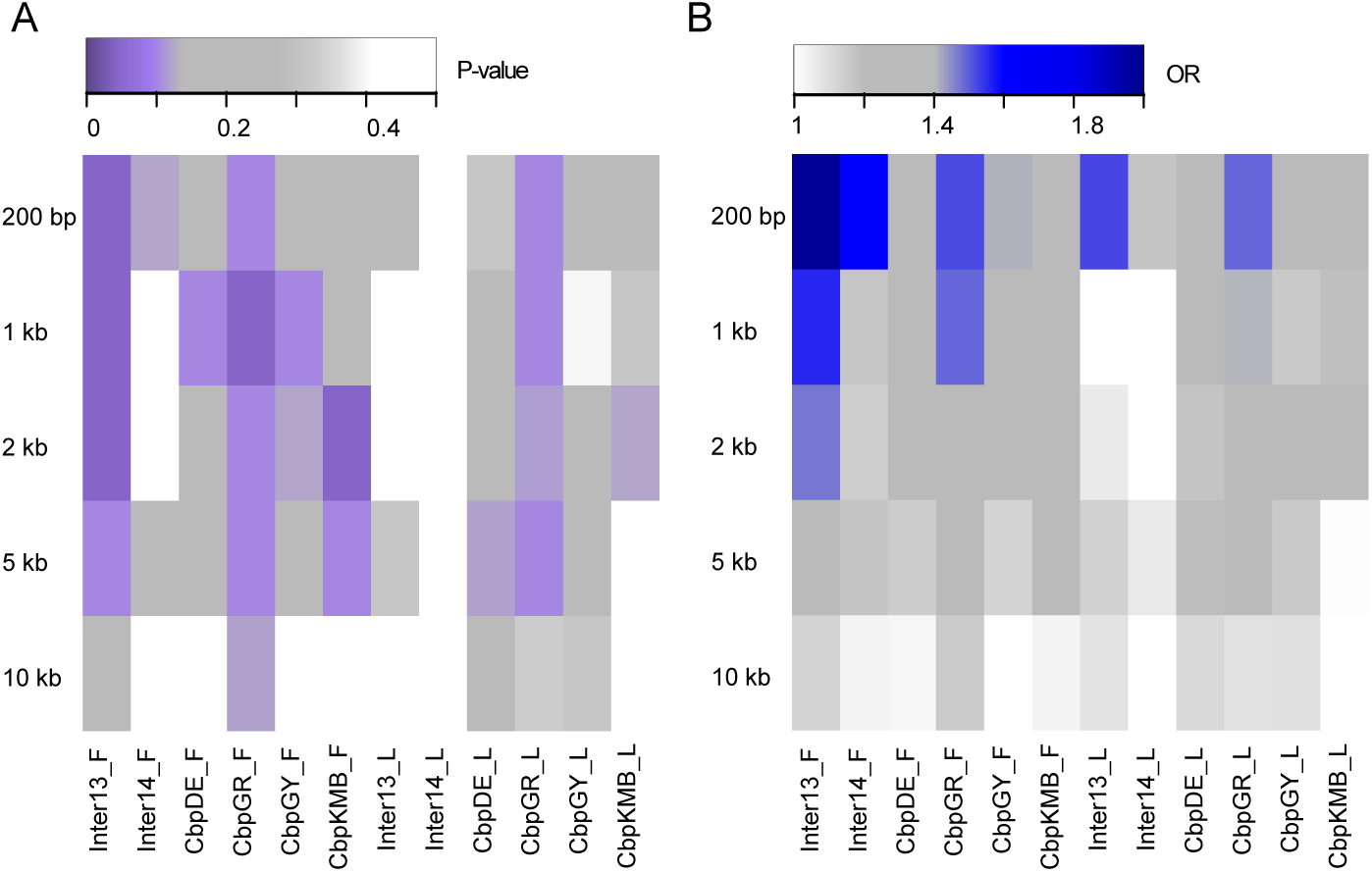
Weak association between heterozygous TEs and expression bias. The heatmaps show multiple-testing corrected P-values (A) and odds ratios (OR) (B) of the association of expression bias and heterozygous TEs within a range of window sizes from 200 bp to 10 kbp from genes. Accession names are as in Table 1 and suffices F and L indicate values for flower buds and leaves, respectively.

### Cis-regulatory divergence among progenitor lineages contributes to homeolog-specific expression

Homeolog-specific expression in *C. bursa-pastoris* has been suggested to be in large part driven by regulatory differences between the progenitor species (Douglas et al 2015). However, this conclusion was based on analysis of differential expression between *C. orientalis* and *C. grandiflora*, and could potentially confound cis-regulatory changes with downstream regulatory effects. Here, we utilized our ASE data to directly assess whether *cis-* regulatory divergence among progenitor lineages are important for homeolog-specific expression in *C. bursa-pastoris*. In agreement with the results of Douglas et al (2015), we find that genes that show a higher expression of the *C. rubella* allele (Cr/Total > 0.5) in the *C. orientalis* x *C. rubella* F1s show a higher expression of the A homeolog in *C. bursa-pastoris* (median CbpA/Total 0.516) in flowers. Genes that show a lower expression of the *C. rubella* allele in the F1s (Cr/Total < 0.5) in turn exhibit a lower expression of the A homeolog in *C. bursa-pastoris* (median CbpA/Total 0.462) in flowers, and comparable patterns were found for leaves (Figure S5). Genes that show a higher expression in the F1s of either the *C. rubella* (Cr/Total > 0.5) or the *C. orientalis* (Cr/Total < 0.5) allele, also show significant expression differences between the *C. bursa-pastoris* homeologs, and this is true for both flower buds (Wilcoxon rank sum test: W = 3864791, p-value < 2.2e-16) and leaves (Wilcoxon rank sum test: W = 3478858, p-value < 2.2e-16). However, it is important to note that many genes, approximately ~35% of those analyzed here, differ in the direction of expression bias among diploid F1s and *C. bursa-pastoris* and thus are not well predicted by current cis-regulatory differences between *C. orientalis* and *C. rubella*.

## Discussion

Here, we have analyzed homeolog-specific expression in the allotetraploid species *C. bursa-pastoris* as well as in diploid F1 hybrids of *C. orientalis* and *C. rubella*, in order to investigate the role of differences in genome-wide TE content for patterns of *cis-*regulatory divergence in association with hybridization and allopolyploidization. Our results demonstrate the potential for variation in patterns of homeolog-specific expression within recently formed tetraploid species such as *C. bursa-pastoris:* while three of four *C. bursa-pastoris* accessions exhibited homeolog expression bias in the direction expected under a model where expression dominance is mediated by TE silencing through the RdDM pathway, the fourth *C. bursa-pastoris* accession showed the opposite direction of homeolog expression bias, as did the diploid F1s. In addition, associations between TE polymorphism and homeolog expression bias were weak and inconsistent among accessions, and the estimated effect of nearby TEs on ASE was small.

Our results are not easily reconciled with a major role for differences in TE content in determining *cis-*regulatory divergence and/or homeolog expression biases in this set of species, and suggest that other factors must be taken into account. These results are therefore similar to those in a recent study of ancient polyploid cotton, where the effect of TEs on gene expression were found to be small (Renny-Byfield et al 2015).

One factor that could be important in explaining our observations is differences in the efficacy of silencing of TEs. In a previous study of *C. grandiflora* x *C. rubella* F1s, we found no evidence for a difference in silencing efficacy, based on the proportion of uniquely mapping 24-nt small RNAs derived from *C. rubella* or *C. grandiflora*, and in that study, there was a stronger positional effect of TEs on ASE (Steige et al 2015a). However, *C. orientalis* and *C. rubella* are substantially more diverged (~1-2 Mya; Douglas et al 2015) than *C. rubella* and *C. grandiflora* (<200 kya; Slotte et al 2013). If *C. orientalis* exhibits more efficient silencing of TEs than *C. rubella*, as might be expected given its smaller genome size (Hurka et al 2012) and lower TE content (Ågren et al 2014) then this might result in preferential silencing of *C. orientalis* alleles, as we observe in the diploid F1s. Differences in silencing efficacy have been observed among closely related species that differ in their mating system, and for instance the selfer *A. thaliana* appears to exhibit a higher efficacy of silencing of TEs than the outcrosser *A. lyrata* (He et al 2011, Hollister et al 2011). Future studies should address this question directly, e.g. using data on uniquely mapping 24-nt small RNA targeting of TEs in these *Capsella* species.

In agreement with Douglas et al (2015), we found that genes that had cis-regulatory differences between *C. orientalis* and *C. rubella* were also more likely to show homeolog-specific expression in the same direction in *C. bursa-pastoris*. Thus, cis-regulatory variation among the progenitors seems to be important for homeolog-specific expression. However, numerous genes deviate from this expectation. Possible explanations for this include epigenetic effects of hybridization and/or polyploidization, or post-polyploidization genetic or epigenetic changes in *C. bursa-pastoris* or the diploid progenitor lineages. Previous studies have found evidence for methylation differences between diploid hybrids and tetraploids of *Brassica* (Ghani et al 2014), and in wheat, polyploidization and hybridization had very different effects on the expression of TE-related small RNAs (Kenan-Eichler et al 2011), which might in turn affect the expression of neighboring genes. Finally, in *Arabidopsis*, changes in ploidy and hybridization affected the expression of siRNAs, and it took several generations to regain stable expression patterns (Ha et al 2009). The extent to which differences in methylation patterns and/or expression of small RNAs might be involved in this case remains unknown.

One caveat to this study is that we assess *cis-*regulatory changes in hybrids between the selfers *C. rubella* and *C. orientalis*, whereas the tetraploid likely originated due to hybridization between an outcrossing ancestor of *C. grandiflora* and *C. rubella* as the pollen parent and a seed parent from the *C. orientalis* lineage (Douglas et al 2015). However, as *C. grandiflora* and *C. rubella* are very closely related (split estimated to have occurred <200 kya; Slotte et al 2013) and phylogenetic trees group either *C. grandiflora* or *C. rubella* as being closest to *C. bursa-pastoris* A (Douglas et al. 2015), using *C. rubella* as one of the hybrid parents should not affect broad patterns strongly. Additionally, we were not able to obtain material for reciprocal crosses of *C. orientalis* and *C. rubella*, but as both our F1s and *C. bursa-pastoris* have *C. orientalis* as the maternal ancestor, this should not affect contrasts of *C. bursa-pastoris* and diploid F1 hybrids.

In sum, the results from this study suggest that differences in TE content alone are not sufficient to explain homeolog-specific expression in *C. bursa-pastoris*, or cis-regulatory changes between *C. orientalis* and *C. rubella*, and that future studies should investigate the role of differences in TE silencing efficacy as well as a broader set of other factors that could affect expression divergence in these species. These results are important for a more general understanding of the underpinnings of *cis-*regulatory divergence in plants.

## Methods

### Plant Material

We included four accessions of *C. bursa-pastoris* from northern and southern Europe and from China. These samples come from the different geographical areas in which separate genetic clusters have been identified in *C. bursa-pastoris* (Slotte et al 2009; Cornille et al 2016) (Supplementary Table S1). In addition, we generated two interspecific diploid hybrids with a genome composition similar to that of *C. bursa-pastoris* by crossing two *C. orientalis* accessions with two accessions of *C. rubella* (all accessions originate from different populations; Table S2).

To avoid accidental self-pollination when crossing *C. orientalis* and *C. rubella*, we emasculated young flower buds before self-fertilization could occur, and hand-pollinated flower buds about three days later. Both crosses had *C. orientalis* as seed parent and *C. rubella* as pollen donor, as reciprocal crosses did not generate viable seeds.

Seeds of all accessions and F1s were surface-sterilized and plated on Murashige-Skoog medium. Plates were vernalized for one week at 4° and germinated in a growth chamber under long day conditions (16 h light, 22°C: 8 h dark, 20°C). Seedlings were moved to soil in pots placed in randomized order in the growth chamber after two weeks. After about 5 weeks, young leaves for RNASeq were collected and flash frozen in liquid nitrogen. About 3 weeks later mixed stages flower buds for RNASeq were collected and flash frozen in liquid nitrogen. We sampled three biological replicates of each F1, consisting of separate F1 individuals from the same cross as well as three biological replicates of each *C. bursa-pastoris* accession. For genomic DNA extraction, we collected leaves of all F1s and their parents as well as all *C. bursa-pastoris* accessions once samples for RNASeq were collected. Ploidy levels were checked by cell flow cytometry of young leaf tissue done by Plant Cytometry Services (Kapel Avezaath Buren, The Netherlands).

### Sample Preparation and Sequencing

We extracted total RNA for whole transcriptome sequencing with the RNEasy Plant Mini Kit (Qiagen, Hilden, Germany), according to the manufacturer’s instructions. For whole genome sequencing, we used a modified CTAB DNA extraction (Doyle and Doyle 1987) to obtain predominantly nuclear DNA. RNA sequencing libraries were prepared using the TruSeq RNA v2 protocol (Illumina, San Diego, CA, USA). DNA sequencing libraries were prepared using the TruSeq DNA v2 protocol. Sequencing was performed on an Illumina HiSeq 2000 instrument (Illumina, San Diego, CA, USA) to gain 100bp paired end reads. Sequencing was done at the Uppsala SNP & SEQ Technology Platform, Uppsala University.

For transcriptome reads, we gained a total of 317.6 Gbp (Q≥30) with an average of 8.8 Gbp per sample (Supplementary Table S3), for genomic reads we a total of 80.2 Gbp (Q≥30) with an average of 8.9 Gbp per sample (Supplementary Table S4). All data generated was uploaded to ENA and can be accessed under project PRJEB12117. The genomic data of Cr39.1 was taken from Steige et al 2015a uploaded to the European Bioinformatics Institute (www.ebi.ac.uk) under PRJEB9020.

### Sequence Quality and Trimming

All DNA and RNA were trimmed using CutAdapt 1.3 (Martin 2011) using custom scripts written by D. G. Scofield. Those scripts identified specific adapters and PCR primers used for each sample and scpecifically removed those. For DNA and RNA Seq reads we removed all read pairs, which had one read shorter than 50 bp. Each sample was individually analysed using fastQC v. 0.10.1 (http://www.bioinformatics.babraham.ac.uk/projects/fastqc/) to identify potential errors that might have occurred during amplification of the DNA or RNA.

### Read Mapping and Variant Calling

Genomic reads were mapped to the *C. rubella* reference genome (v 1.0) using BWA-MEM (Li 2013) with default parameters. Variant calling was done using GATK v.3.3.0 (McKenna et al. 2010) according to GATK best practices (DePristo et al. 2011, Van der Auwera et al. 2013). In brief, this includes steps to mark duplicates, doing realignment around indels and recalibrate base quality scores using a set of 1.3 million known SNPs in *C. grandiflora* (Williamson et al. 2014) as known variants. Only SNPs with high quality were kept for further analyses. Variant discovery was done jointly for the different parental accessions of the F1s using UnifiedGenotyper.

### Reconstruction of parental haplotypes of interspecific FIs

To reconstruct parental haplotypes of interspecific F1s, we used genomic data of the parental accessions. We did read mapping and variant calling of the parental genomic reads as described above. The vcf files gained from this were used together with the *C. rubella* reference genome to generate new reference genomes, containing the specific genome-wide haplotypes of the F1s using custom java scripts by Johan Reimegård. Afterwards, read mapping of both transcriptomic and genomic reads were done against the specific parental haplotypes using STAR v.2.3.0.1 (Dobin et al. 2013) and read counts at all reliable SNPs (see “Filtering”) were obtained using Samtools mpileup and a custom software written in javascript by Johan Reimegård. For *C. bursa-*pastoris all mapping was done to the parental haplotypes reconstructed for *C. orientalis* x *C. rubella* F1 Inter13, and sites were subsequently filtered as described in the section Filtering below. The files containing genomic and transcriptomic allele counts were used to assess allele specific biases in the F1s and homeolog expression biases in the *C. bursa-pastoris* accessions.

### Filtering

To retain genomic regions where we have high confidence in our SNP calls, we removed genomic regions that show an elevated fraction of repeats and selfish genetic elements, as in Steige et al 2015a. The bedfile containing these regions were taken from phytozome (http://phytozome.jgi.doe.gov/pz/portal.html) and were included in the publication of the *C. rubella* reference genome (Slotte et al. 2013). Additionally we removed regions that had unusually high proportions of heterozygous calls in an inbred *C. rubella* line, which was assessed in a previous publication (Steige et al. 2015a). To only retain SNPs that are informative about differences between the parental species (and therefore the two homeologs) of *C. bursa-pastoris*, we only kept sites that had a fixed difference between a set of 12 scattered *C. grandiflora* (Steige et al 2015b; Hatorangan MR, Laenen B, Steige K, Slotte T, Köhler C, submitted) and 10 *C. orientalis* accessions and showed fixed heterozygosity within *C. bursa-pastoris*, similar to the filtering conducted in Douglas et al (2015).

### Analysis of allele-specific expression

Analyses of allele-specific expression (ASE) and homeolog-specific expression (HSE) were done as described in Steige et al 2015a. In short, we used a hierarchical Bayesian method developed by Skelly et al 2011, which has a reduced rate of false positives and naturally incorporates replicates in the analysis. The method uses genomic data to fit parameters to a beta-binomial distribution of variation in allelic ratios due to technical variation (as there is no true allelic bias in genomic data). These parameters are then used in the analysis of the RNA reads, which assigns a posterior probability of a gene showing ASE.

### Identifying insertions of transposable elements (TEs)

We identified insertions of transposable elements in our F1s as well as the four *C. bursa-pastoris* accessions as was described in (Steige et al. 2015a). In short, we used the genomic data to infer TE insertions using the PoPoolationTE pipeline (Kofler et al. 2012), modified to require a minimum of 5 reads supporting a TE, and using TE sequences from a library based on several Brassicaceae species (Slotte et al. 2013). We inferred presence of heterozygous TEs and homozygous TEs by their frequency as in Ågren et al (2014) and Steige et al (2015a). We tested for enrichment of TEs close by genes showing ASE or HSE using a Fisher exact tests, and a range of window sizes for scoring TE insertions near genes (200 bp, 1 kbp, 2 kbp, 5k bp, 10kbp). P-values were corrected for multiple testing using the Benjamini and Hochberg method (Benjamini and Hochberg 1995).

## Acknowledgments

We thank Barbara Neuffer for kindly providing seeds of *C. orientalis*. This study was funded by a grant from the Swedish Research Council (to T.S.) and a European Research Council Starting Independent Researcher grant (to C.K.).

## Literature Cited

Alexa A, Rahnenführer J, Lengauer T. 2006. Improved scoring of functional groups from gene expression data by decorrelating GO graph structure. Bioinformatics 22:1600–1607.

Barker MS, Arrigo N, Baniaga AE, Li Z, Levin DA. 2015. On the relative abundance of autopolyploids and allopolyploids. New Phytologist doi: 10.1111/nph.13698

Benjamini Y, Hochberg Y. 1995. Controlling the False Discovery Rate: A Practical and Powerful Approach to Multiple Testing. J R Statist Soc B 57:289–300.

Chater AO. 1993. Capsella. In: Tutin TG, Heywood H, Burges NA, Moore DM, Valentine DH, Walters SM, Webb DA, editors. Flora Europaea. Cambridge, UK: Cambridge University Press. pp. 381–382.

Cornille A, Salcedo A, Kryvokhyza D, Glémin S, Holm K, Lascoux M. 2016. Genomic signature of successful colonization of Eurasia by the allopolyploid shepherd's purse (Capsella bursa-pastoris). Molecular Ecology 25: 616–629.

DePristo MA, Banks E, Poplin R, Garimella KV, Maguire JR, Hartl C, Philippakis AA, del Angel G, Rivas MA, Hanna M, McKenna A, Fennell TJ, Kernytsky AM, Sivachenko AY, Cibulskis K, Gabriel SB, Altshuler D, Daly MJ. 2011. A framework for variation discovery and genotyping using next-generation DNA sequencing data. Nat Genet 43:491–498.

Dobin A, Davis CA, Schlesinger F, Drenkow J, Zaleski C, Jha S, Batut P, Chaisson M, Gingeras TR. 2013. STAR: ultrafast universal RNA-seq aligner. Bioinformatics 29:15–21.

Douglas GM, Gos G, Steige KA, Salcedo A, Holm K, Josephs EB, Arunkumar R, Ågren JA, Hazzouri KH, Wang W, Platts AE, Williamson RJ, Neuffer B, Lascoux M, Slotte T, Wright SI. 2015. Hybrid origins and the earliest stages of diploidization in the highly successful recent polyploid Capsella bursa-pastoris. Proceedings of the National Academy of Sciences. 112:2806–2811

Doyle JJ, Doyle JL. 1987. A rapid DNA isolation procedure for small quantities of fresh leaf tissue. Phytochem bull 19: 11–15.

Flagel LE, Wendel JF. 2010. Evolutionary rate variation, genomic dominance and duplicate gene expression evolution during allotetraploid cotton speciation. New Phytol 186:184–193.

Freeling M, Woodhouse MR, Subramaniam S, Turco G, Lisch D, Schnable JC. 2012. Fractionation mutagenesis and similar consequences of mechanisms removing dispensable or less-expressed DNA in plants. Curr Opin Plant Biol 15:131–139.

Ghani MA, Li J, Rao L, Raza MA, Cao L, Yu N, Zou X, Chen L. 2014. The role of small RNAs in wide hybridisation and allopolyploidisation between Brassica rapa and Brassica nigra. BMC Plant Biology 14:272.

Ha M, Lu J, Tian L, Ramachandran V, Kasschau KD, Chapman EJ, Carrington JC, Chen X, Wang X-J, Chen ZJ. 2009. Small RNAs serve as a genetic buffer against genomic shock in Arabidopsis interspecific hybrids and allopolyploids. Proceedings of the National Academy of Sciences 106:17835–17840.

He F, Zhang X, Hu J-Y, Turck F, Dong X, Goebel U, Borevitz JO, de Meaux J. 2012. Widespread interspecific divergence in cis-regulation of transposable elements in the Arabidopsis genus. Mol Biol Evol 29:1081–1091.

Hollister JD, Gaut BS. 2009. Epigenetic silencing of transposable elements: a trade-off between reduced transposition and deleterious effects on neighboring gene expression. Genome Res 19:1419–1428.

Hollister JD, Smith LM, Guo Y-L, Ott F, Weigel D, Gaut BS. 2011. Transposable elements and small RNAs contribute to gene expression divergence between Arabidopsis thaliana and Arabidopsis lyrata. Proceedings of the National Academy of Sciences 108:2322–2327.

Hurka H, Neuffer B. 1997. Evolutionary processes in the genus Capsella (Brassicaceae). Plant Syst Evol 206:295–316.

Hurka H, Friesen N, German DA, Franzke A, Neuffer B. 2012. ‘Missing link’ species Capsella orientalis and Capsella thracica elucidate evolution of model plant genus Capsella (Brassicaceae). Mol Ecol doi: 10.1111/j.1365–294X.2012.05460.x

Jalas J, Suominen J, eds. 1994. Atlas Florae Europaeae. Distribution of Vascular Plants in Europe. Vol 10. Cruciferae (Sisymbrium to Aubrieta). Helsinki, Finland: The Committee for Mapping the Flora of Europe & Societas Biologica Fennica Vanamo.

Kenan-Eichler M, Leshkowitz D, Tal L, Noor E, Melamed-Bessudo C, Feldman M, Levy AA. 2011. Wheat hybridization and polyploidization results in deregulation of small RNAs. Genetics 188:263–272.

Kofler R, Betancourt AJ, Schlötterer C. 2012. Sequencing of pooled DNA samples (Pool-Seq) uncovers complex dynamics of transposable element insertions in Drosophila melanogaster. PLoS Genet 8:e1002487.

Langham RJ, Walsh J, Dunn M, Ko C, Goff SA, Freeling M. 2004. Genomic duplication, fractionation and the origin of regulatory novelty. Genetics 166:935–945.

Li A, Liu D, Wu J, Zhao X, Hao M, Geng S, Yan J, Jiang X, Zhang L, Wu J, Yin L, Zhang R, Wu L, Zheng Y, Mao L. 2014. mRNA and Small RNA Transcriptomes Reveal Insights into Dynamic Homoeolog Regulation of Allopolyploid Heterosis in Nascent Hexaploid Wheat. Plant Cell 26:1878–1900.

Li H. 2013. Aligning sequence reads, clone sequences and assembly contigs with BWA-MEM. arXiv:1303.3997

Lippman Z, Gendrel A-V, Black M, Vaughn MW, Dedhia N, McCombie WR, Lavine K, Mittal V, May B, Kasschau KD, Carrington JC, Doerge RW, Colot V, Martienssen R. 2014. Role of transposable elements in heterochromatin and epigenetic control. Nature 430:471–476.

Martin M. 2011. Cutadapt removes adapter sequences from high-throughput sequencing reads. EMBnet.journal. 17:10–12.

Matzke MA, Mosher RA. 2014. RNA-directed DNA methylation: an epigenetic pathway of increasing complexity. Nat Rev Genet 15:394–408.

McKenna A, Hanna M, Banks E, Sivachenko A, Cibulskis K, Kernytsky A, Garimella K, Altshuler D, Gabriel S, Daly M, DePristo MA. 2010. The Genome Analysis Toolkit: a MapReduce framework for analyzing next-generation DNA sequencing data. Genome Res. 20:1297–1303.

Ramsey J, Schemske D. 1998. Pathways, mechanisms, and rates of polyploid formation in flowering plants. Annu Rev Ecol Syst 29:467–501.

Renny-Byfield S, Gong L, Gallagher JP, Wendel JF. 2015. Persistence of subgenomes in paleopolyploid cotton after 60 my of evolution. Mol Biol Evol 32:1063–1071.

Schnable JC, Springer NM, Freeling M. 2011. Differentiation of the maize subgenomes by genome dominance and both ancient and ongoing gene loss. Proceedings of the National Academy of Sciences 108:4069–4074.

Skelly DA, Johansson M, Madeoy J, Wakefield J, Akey JM. 2011. A powerful and flexible statistical framework for testing hypotheses of allele-specific gene expression from RNA-seq data. Genome Res. 21:1728–1737.

Slotte T, Hazzouri KM, Agren JA, Koenig D, Maumus F, Guo YL, Steige K, Platts AE, Escobar JS, Newman LK, Wang W, Mandáková T, Vello E, Smith LM, Henz SR, Steffen J, Takuno S, Brandvain Y, Coop G, Andolfatto P, Hu TT, Blanchette M, Clark RM, Quesneville H, Nordborg M, Gaut BS, Lysak MA, Jenkins J, Grimwood J, Chapman J, Prochnik S, Shu S, Rokhsar D, Schmutz J, Weigel D, Wright SI. 2013. The Capsella rubella genome and the genomic consequences of rapid mating system evolution. Nat Genet. 2013 45:831–835.

Slotte T, Huang HR, Holm K, Ceplitis A, St. Onge K, Chen J, Lagercrantz U, Lascoux M. 2009. Splicing variation at a FLOWERING LOCUS C homeolog is associated with flowering time variation in the tetraploid Capsella bursa-pastoris. Genetics 183:337–345.

Soltis DE, Albert VA, Leebens-Mack J, Bell CD, Paterson AH, Zheng C, Sankoff D, dePamphilis CW, Wall PK, Soltis PS. 2009. Polyploid and angiosperm diversification. American Journal of Botany 96:336–348.

Steige KA, Reimegård J, Koenig D, Scofield DG, Slotte T. 2015a. Cis-Regulatory Changes Associated with a Recent Mating System Shift and Floral Adaptation in Capsella. Mol Biol Evol 32:2501–2514.

Steige KA, Laenen B, Reimegård J, Scofield DG, Slotte T. 2015b. The impact of natural selection on the distribution of cis-regulatory variation across the genome of an outcrossing plant. bioRxiv 10.1101/034025.

Steige KA, Slotte T. 2016. Genomic legacies of the progenitors and the evolutionary consequences of allopolyploidy. Curr Opin Plant Biol 30:88–93.

Van der Auwera GA, Carneiro MO, Hartl C, Poplin R, Del Angel G, Levy-Moonshine A, Jordan T, Shakir K, Roazen D, Thibault J, Banks E, Garimella KV, Altshuler D, Gabriel S, DePristo MA. 2013. From FastQ data to high confidence variant calls: the Genome Analysis Toolkit best practices pipeline. Curr Protoc Bioinformatics 11:11.10.1–11.10.33.

Wang X, Weigel D, Smith LM. 2013. Transposon variants and their effects on gene expression in Arabidopsis. PLoS Genet. 9:e1003255.

Williamson RJ, Josephs EB, Platts AE, Hazzouri KM, Haudry A, Blanchette M, Wright SI. 2014. Evidence for widespread positive and negative selection in coding and conserved noncoding regions of Capsella grandiflora. PLoS Genet 10:e1004622.

Wood TE, Takebayashi N, Barker MS, Mayrose I, Greenspoon PB, Rieseberg LH. 2009. The frequency of polyploid speciation in vascular plants. Proceedings of the National Academy of Sciences 106:13875–13879.

Woodhouse MR, Cheng F, Pires JC, Lisch D, Freeling M, Wang X. 2014. Origin, inheritance, and gene regulatory consequences of genome dominance in polyploids. Proceedings of the National Academy of Sciences 111:5283–5288.

Ågren JA, Huang H-R, Wright SI. 2016. Transposable element evolution in the allotetraploid Capsella bursa-pastoris and the perfect storm hypothesis. BioRxiv doi:10.1101/042325

Ågren JA, Wang W, Koenig D, Neuffer B, Weigel D, Wright SI. 2014. Mating system shifts and transposable element evolution in the plant genus Capsella. BMC Genomics. 15:602.

